# Identification of neoantigens in esophageal adenocarcinoma

**DOI:** 10.1101/2022.05.04.490567

**Authors:** Ben Nicholas, Alistair Bailey, Katy J McCann, Oliver Wood, Robert C Walker, Robert Parker, Nicola Ternette, Tim Elliott, Tim J Underwood, Peter Johnson, Paul Skipp

**Affiliations:** Centre for Proteomic Research, Biological Sciences and Institute for Life Sciences, Building 85, University of Southampton, SO17 1BJ, UK; Centre for Cancer Immunology and Institute for Life Sciences, Faculty of Medicine, University of Southampton, UK; School of Cancer Sciences, Faculty of Medicine, University of Southampton, Southampton, UK; Centre for Cellular and Molecular Physiology, Nuffield Department of Medicine, University of Oxford, Oxford, UK; Centre for Immuno-oncology, Nuffield Department of Medicine, University of Oxford, UK; Cancer Research UK Clinical Centre, University of Southampton, Southampton, UK

**Keywords:** HLA, peptidome, esophageal adenocarcinoma, antigen presentation

## Abstract

Esophageal adenocarcinoma (EAC) has a relatively poor long-term survival and limited treatment options. Promising targets for immunotherapy are short peptide neoantigens containing tumor mutations, presented to cytotoxic T-cells by human leukocyte antigen molecules (HLA). Despite an association between putative neoantigen abundance and therapeutic response across cancers, immunogenic neoantigens are challenging to identify. Here we characterized the mutational and immunopeptidomic landscapes of tumors from a cohort of seven patients with EAC. We directly identified one HLA-I presented neoantigen from one patient, and report functional T-cell responses from a predicted HLA-II neoantigen in a second patient. The predicted class II neoantigen contains both HLA I and II binding motifs. Our exploratory observations are consistent with previous neoantigen studies in finding that neoantigens are rarely directly observed, and an identification success rate following prediction in the order of 10%. However, our identified putative neoantigen is capable of eliciting strong T-cell responses, emphasizing the need for improved strategies for neoantigen identification.

## Introduction

Esophageal adenocarcinoma (EAC) is the 14th most common cancer in the UK, with a 10-year survivability of 12% [1]. Early-stage treatment of EAC involves resection of the esophagus, whereas later stage diagnosis is treated with chemoradiotherapy or chemotherapy followed by surgery [2]. Relative to other cancers, EAC is characterized by having a high mutational burden, measured as the number of mutations per protein coding region [3]. Many of these mutations appearing in EAC driver genes [4,5].

tumor infiltrating lymphocytes (TILs), specifically cytotoxic CD8+ and CD4+ helper T-cells recognize respectively, peptides of intracellular and extracellular origin presented by class I and II human leukocyte molecules (HLA). Presented at the cell surface, these HLA bound peptides form the immunopeptidome. Neoantigen peptides contain tumor mutations, making attractive therapeutic targets because of their potential to elicit tumor specific T-cell responses.

Progress in developing neoantigen vaccines has been hindered by the difficulty in identifying neoantigen targets, and the challenge of overcoming the immunosuppressive tumor microenviroment. In addressing neoantigen identification, direct identification using immunopeptidomics suggests observing neoantigens is rare [6]. Attempts to predict neoantigens using HLA binding algorithms show it is relatively straight forward to create a long list of potential putative neoantigens, but difficult to reliably select immunogenic neoantigens [7]. Large scale studies of EAC report the density of CD8+ T-cells correlates with the number of somatic mutations [4], but an analysis of the EAC immunopeptidome has yet to be performed.

Here we explore a proteogenomics approach combining whole exome sequencing (WES), gene expression (RNASeq), HLA immunopeptidomics and algorithmic neoantigen prediction to identify neoantigens in seven EAC patients. We show that EAC has an abundance of somatic mutations and immunopeptides, and that whilst direct observation or prediction of immunogenic neoantigens remains challenging we were able to identify two neoantigens in two patients, one by direct observation and one by prediction. These findings are an important step towards demonstrating the usefulness of neoantigen based therapies for EAC.

## Results

We collected tissues comprising of tumor and adjacent normal tissue, and peripheral blood mononuclear cells (PBMCs) from seven male individuals with esophageal adenocarcinoma (median age 68; Table 1). Whole genome sequencing for three donors have been previously deposited as part of ICGC project ESAD-UK and EGA data set EGAD00001007785. We sequenced the exomes of tumor and normal tissues, and performed gene expression and immunopeptidomic analysis of the tumor tissues. PBMCs were used for HLA typing and I FN-γ ELISpot functional assays for patient EN-181-11 (Figure 1) [8].

**Figure 1:**
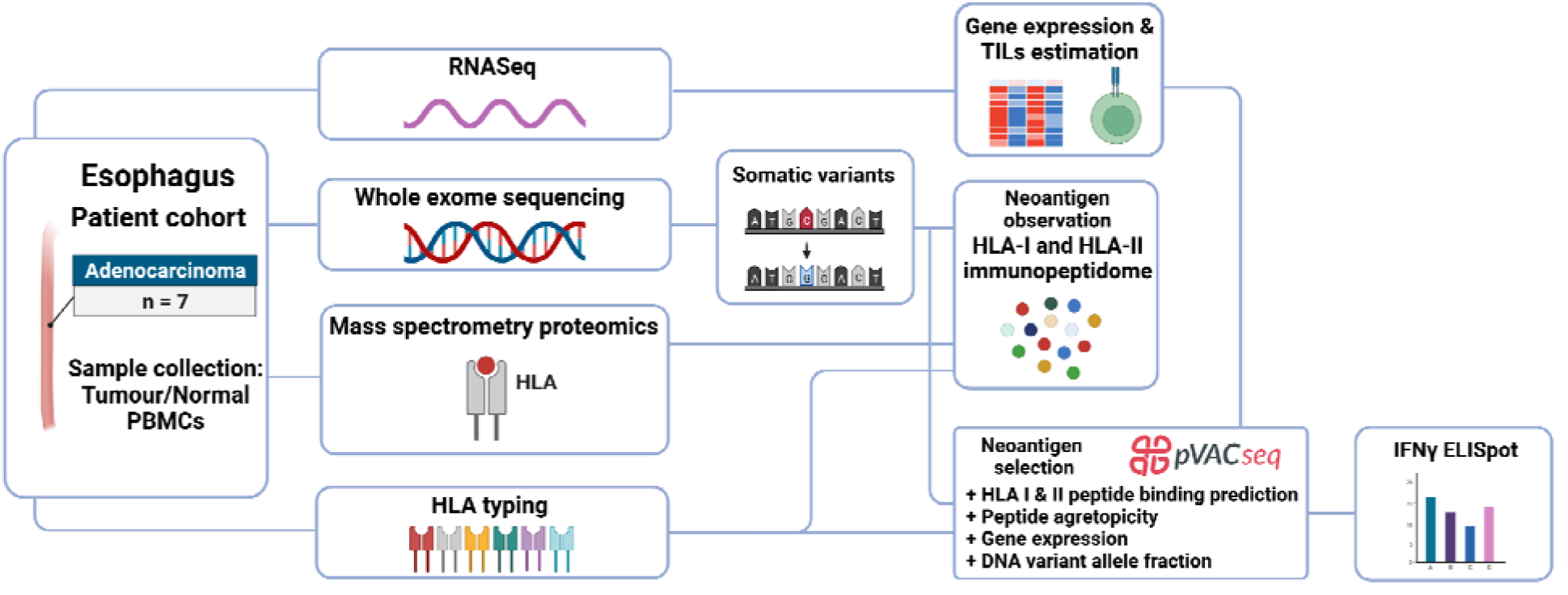
Workflow of the approach to identify HLA-I and -II neoantigens isolated from
 esophageal tissues.

**Table 1:** Clinical summary of patients in this study with esophageal adenocarcinoma

| Donor | ICGC Donor | Age at diagnosis | Sex | Tumor location | Tumor stage | Treatment modality |
| --- | --- | --- | --- | --- | --- | --- |
| EN-181-11 | DO234382 | 64 | Male | Gastro-Esophageal junction (Siewert II) | 2 | Surgery |
| EN-216-11 | DO50307 | 74 | Male | Lower Esophagus | 2 | Surgery |
| EN-430-11 |  | 78 | Male | Lower Esophagus | 3 | Chemotherapy + Surgery |
| EN-454-11 | DO50387 | 81 | Male | Gastro-Esophageal junction (Siewert II) | 3 | Surgery |
| EN-489-12 |  | 62 | Male | Lower Esophagus | 2 | Chemotherapy + Surgery |
| EN-711-16 |  | 68 | Male | Lower Esophagus | 3 | Chemotherapy + Surgery |
| EN-716-11 |  | 66 | Male | Lower Esophagus | 3 | Chemotherapy + Surgery |

### The mutational landscape of seven esophageal adenocarcinomas

To assess the likelihood of identifying HLA presented neoantigens we first examined the mutational landscape of the seven esophageal adenocarcinomas. Somatic mutations accumulate in the genome over time as cells divide, and in cancer the causes and patterns of somatic mutations help characterize the cancer type and explain its cells selective advantage [3]. The total number of somatic mutations per coding region of genome defines the mutational burden of the cancer type, and has been correlated with response to anti-PD1 therapy, and is therefore a proxy for the number of neoantigens presented by tumor cells [9,10]. Across cancers, median mutational burden ranges from 16 to over 300 mutations per megabase [3]. Here, the esophageal adenocarcinomas had a median mutational burden of 124 mutations/Mb in comparison to a median of 40 mutations/Mb for normal adjacent tissue (Figure 2A, S1 Table). Four patterns of single base substitutions created by the somatic mutations were extracted as mutational signatures and fitted to those identified in COSMIC [11–13] (Figure 2B-F). Different fractions of these four signatures were seen in each individual (Figure 2B). Extracted signatures A and B contain high proportions of C>T substitutions, fitting signatures SBS1 and SBS5 respectively (Figure 2C,F). These are both clock like signatures correlated with ageing and have previously been reported in large scale studies of EAC [4,12]. Extracted signatures C and D fit signatures SBS43 and SBS55 respectively, and contain high proportions of T>G substitutions (Figure 2A,C). COSMIC flags these signatures as possible sequencing artefacts, but high proportions of T>G substitutions have been reported as characteristic of EAC, and indicative of high levels of neoantigen presentation [4].

**Figure 2:**
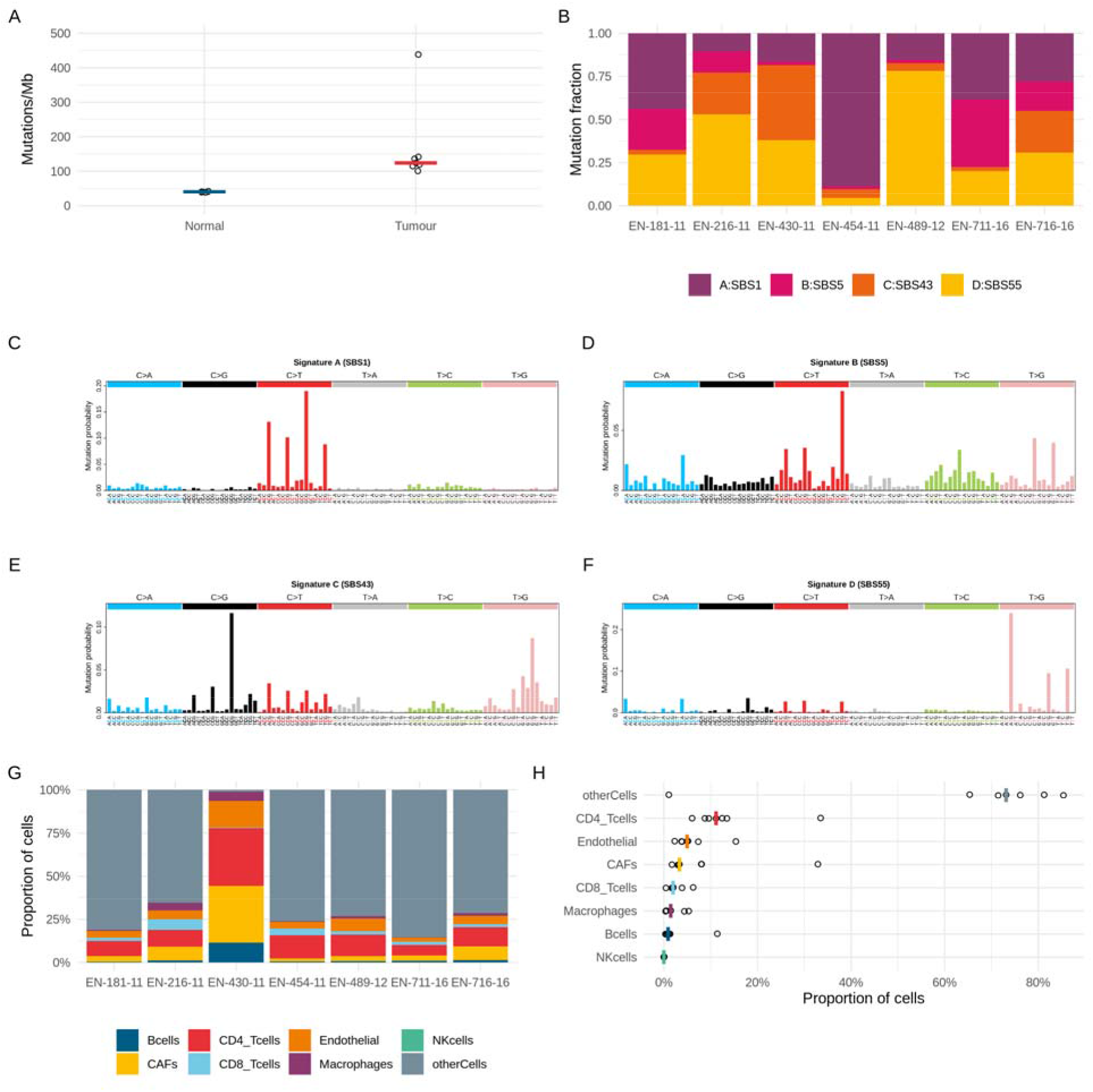
The mutational landscape of seven esophageal adenocarcinomas (A) The mutational burden of tumor and normal adjacent tissues from WES assuming a whole exome size of 30 Mb. The bar demarks the median. (B) The proportions of the four single base substitution mutational signatures in each EAC sample extracted from WES. The best fit signatures in COSMIC v3 database are prefixed SBS. (C-F) The four mutational signatures extracted from WES of seven EAC samples. (G) The proportions of TILs estimated from bulk tumor RNASeq in each EAC sample. (H) The proportions of TILs estimated from bulk tumor RNASeq across the cohort. The bar demarks the median.

The immune response to neoantigens is also contingent on the ability of immune cells to infiltrate tumors. Using bulk tumor gene expression data, we estimated the fraction of tumor infiltrating lymphocytes present in our tumors [14] (Figure 2G-H). This estimation also indicates a measure of tumor purity by collating all the genes not corresponding with TILs as ‘otherCells’, which we would expect to make up the majority of the cells in a tumor sample. Therefore the estimation of only 1% other cells for EN-430-11 as compared with a median of 73% indicates that the biopsy captured predominantly non-tumor material. The remaining six samples have similar proportions of CD4+ and CD8+ TILs, with median fractions of 11% and 2% respectively, indicating the presence of TILs, a necessary but not sufficient requirement for a response to presented neoantigens.

In summary, the mutational landscape of our EAC samples is characterized by a high tumor mutational burden along with the presence of TILs, both necessary conditions for neoantigen presentation and recognition respectively.

### Immunopeptidome analysis reveals one putative neoantigen

We next sought to directly observe neoantigens present in the immunopeptidomes of our EAC samples. Using the mutations identified from WES we created individual databases appended with patient specific mutated sequences (mutanomes) to search for neoantigens in their immunopeptidomes (Figure 1). In total we identified 41,535 HLA class I and II peptides isolated from these tumors by LC-MS/MS analysis at a false discovery rate of 1% (S1 Table). These immunopeptidomes comprised of 24,095 unique class I and 8,023 unique class II peptides, forming characteristic HLA length distributions with modes of 9-mers and 15-mers for HLA class I and II peptides respectively (Figure 3A).

**Figure 3:**
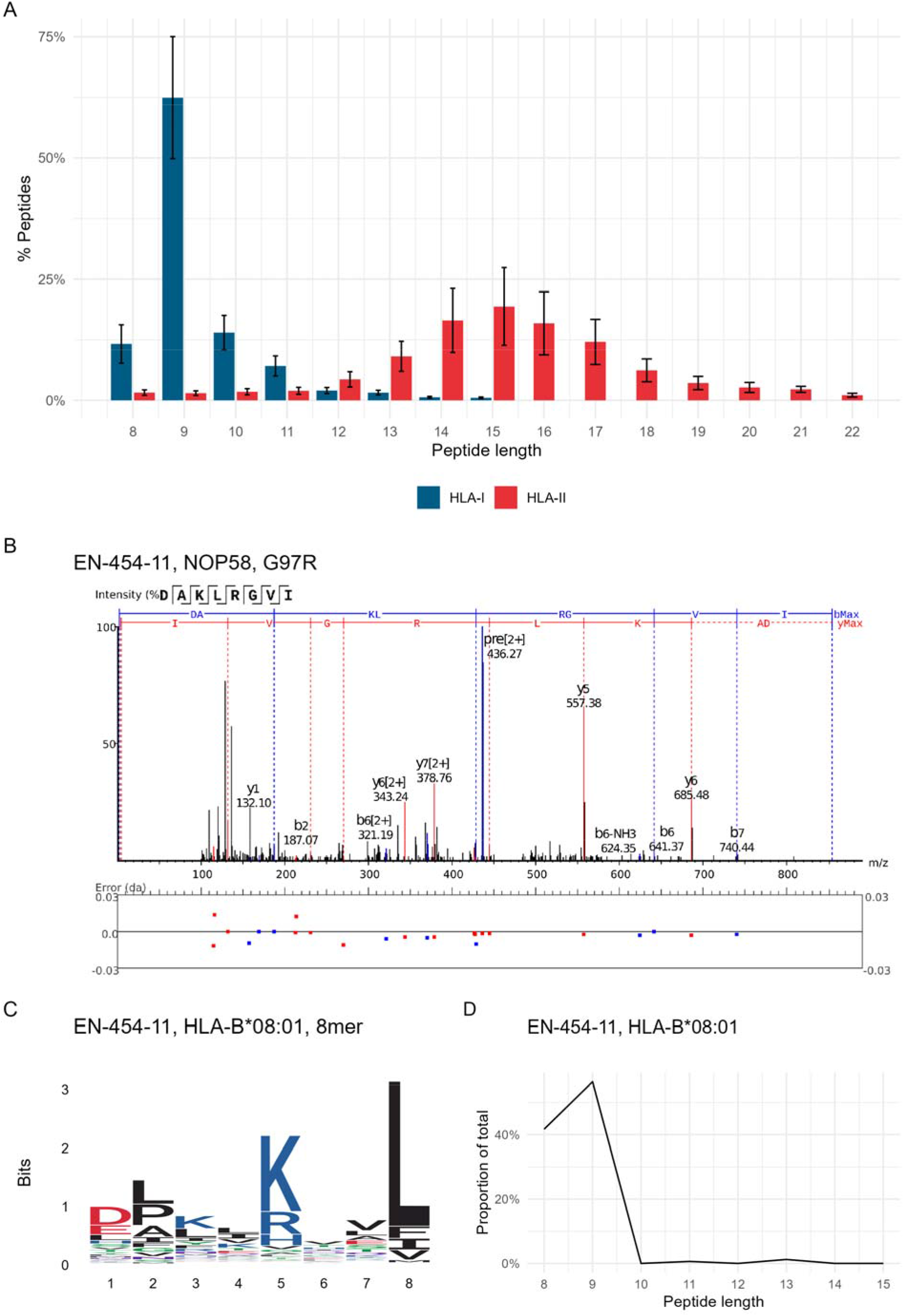
A single putative neoantigen identifed from the immunopeptidomes of seven esophageal adenocarcinomas (A) Histogram of 41,535 eluted HLA class I and II peptides from seven EAC samples. (B) MS/MS spectrum from donor EN-454-11 of putative HLA-B*08:01 8-mer neoantigen DAKLRGVI. (C) Motif of all 8-mer peptides from donor EN-454-11 assigned to HLA-B*08:01. (D) Length distribution of n=650 peptides from donor EN-454-11 assigned to HLA-B*08:01.

Across the seven patients we identified only one putative neoantigen from the HLA-I immunopeptidome of patient EN-454-11 (Figure 3B) [15]. This is an 8-mer peptide derived from Nucleolar protein 58 (Gene: NOP58, UniProt: Q9Y2X3 COSMIC: COSV51895876) with a hydrophobic glycine to basic arginine mutation at protein amino acid (AA) residue 97, peptide AA residue 5. The mutation at peptide residue 5 creates a sequence DAKLRGVI with anchor residues for HLA-B*08:01 (Figure 3C) [16,17]. Over 40% of the 650 HLA-B*08:01 peptides identified for EN-454-11 were 8-mers, consistent with previous reports of a secondary length preference for 8-mers for this allotype (Figure 3D) [17]. Unfortunately, there were insufficient PBMCs available to perform functional assays for this donor, so we next focused on predicted neoantigens from patient EN-181-11 from which we could perform a functional assay.

### Neoantigen prediction and functional analysis identifies a neoantigen with both class I and II HLA motifs

Neoantigen predictions from the EN-181-11 mutanome of 8-11mer peptides for class I HLA-A and B, and of 15-mer peptides for class II DRB1 allotypes were calculated using pVACseq [18,19]. Neoantigen rank score is calculated as a function of the predicted binding affinity, the neoantigen agretopicity (the relative increase in predicted binding affinity of mutant peptide to wildtype peptide), the variant allele frequency and gene expression levels (Figure 1, Materials and Methods). Predictions were performed for each peptide length and allotype combination yielding 15 ranked tables, comprising a total of 6842 peptides with binding affinity <500 nM for patient EN-181-11. Nine top ranking putative neoantigens were selected for functional analysis (Table 2, Supporting Information).

**Table 2:**
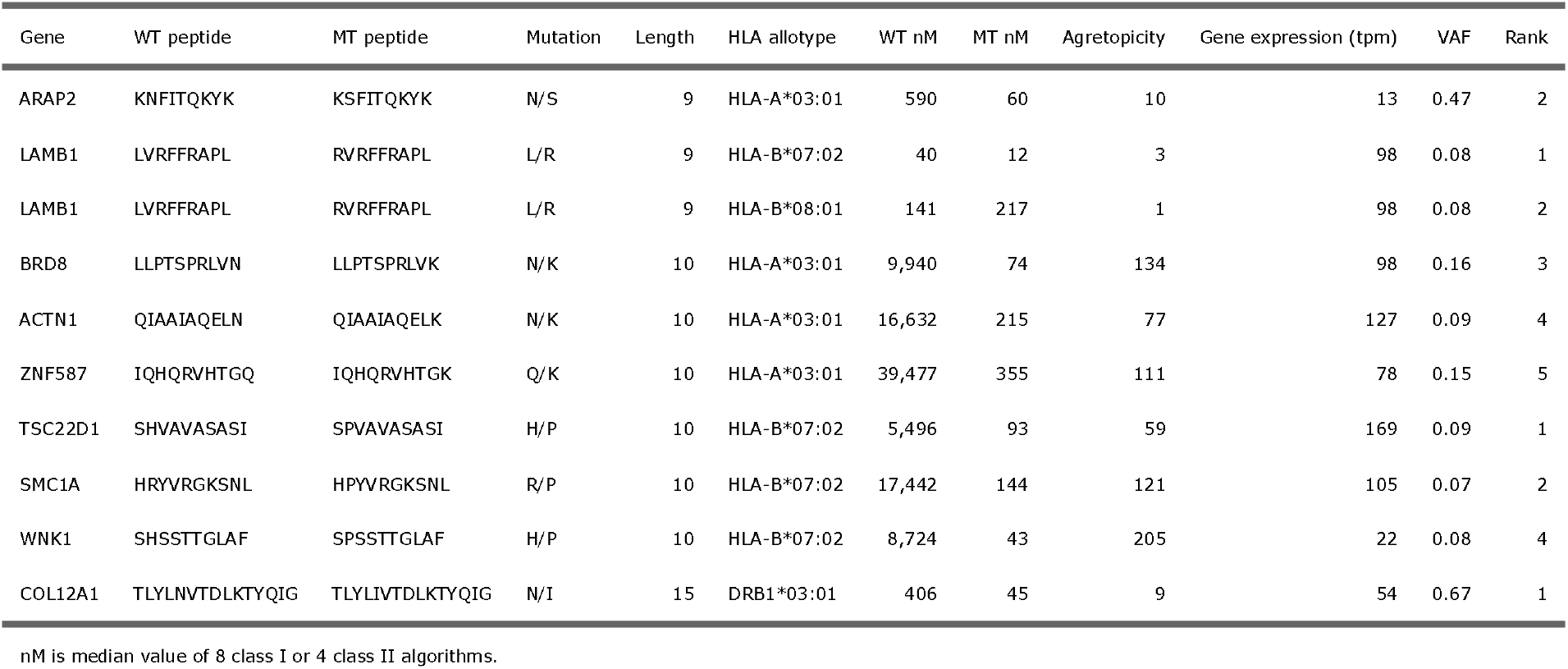
Summary of top EN-181-11 predicted neoantigens

We synthesized both the neoantigen and wild type peptides at their specific lengths (Table 2) and tested their ability to stimulate T-cells present in PBMCs using an IFN-□ release cultured ELISpot assay (Figure 4A). We observed a strong response for only the putative class II neoantigen derived from collagen alpha-1(XII) chain (Gene: COL12A1 UniProt: Q99715). Closer examination of the COL12A1 neoantigen sequence revealed that the first nine amino acids TLYLIVTDLK contain the HLA-A*03:01 motif in addition to the HLA-DRB1*03:01 motif in the full length TLYLIVTDLKTYQIG peptide (Figure 4B-C). Moreover, the observation of COL12A1 wild type peptides in both the class I and II immunopeptidomes of EN-181-11 indicate that this protein is presented in both antigen processing pathways by this tumor (Supporting Information).

**Figure 4:**
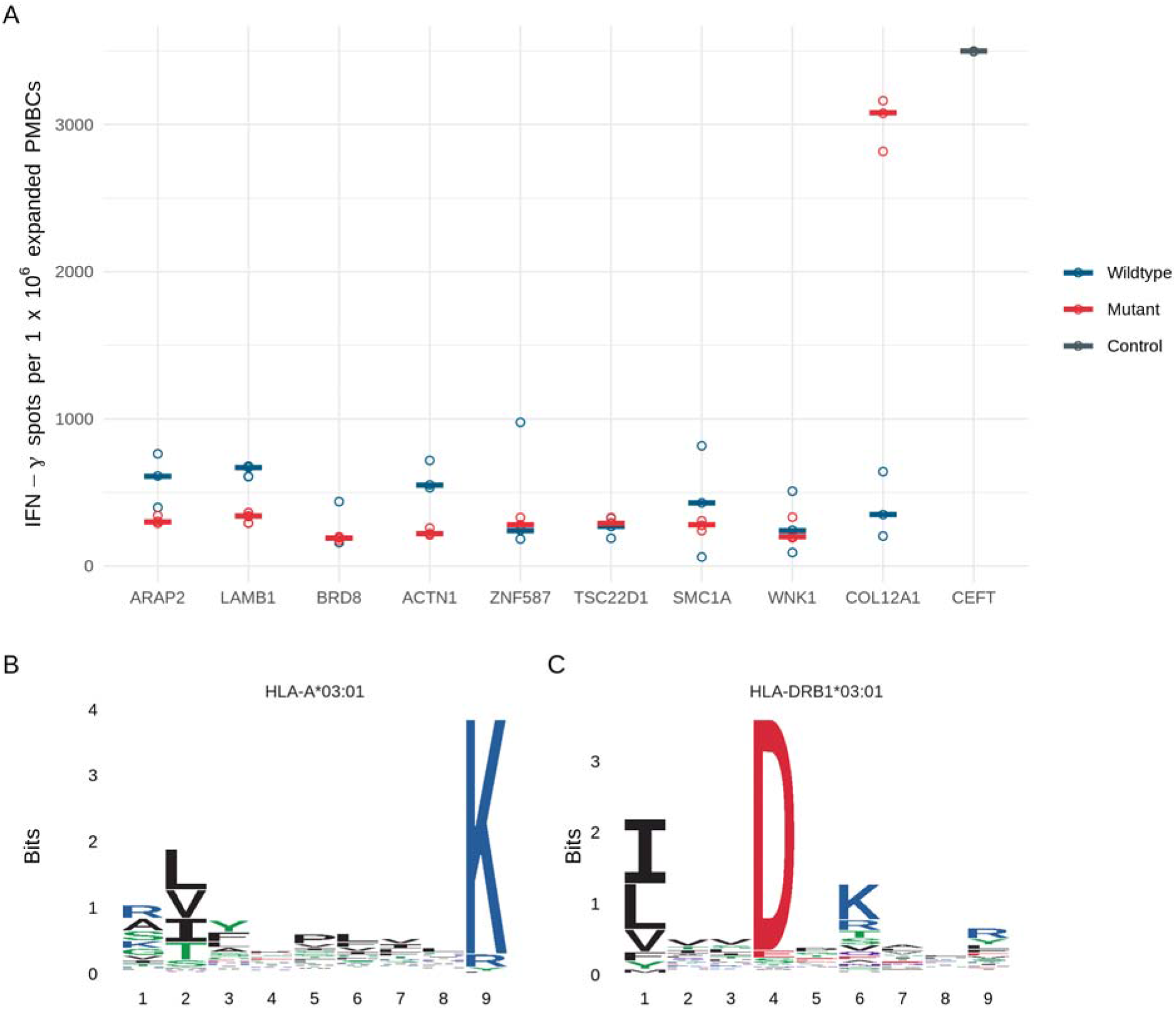
Functional T-cell assay identifies a responding neoantigen containing class I and II HLA motifs (A) IFN-γ ELISpot of nine predicted neoantigen peptides (mutant) and their wildtype equivalents. Lengths and sequences as provided in Table 2. (B) HLA-A*03:01 9-mer motif assigned to EN-181-11 observed peptides (C) HLA-DRB1*03:01 core motif assigned to EN-181-11 observed peptides

## Discussion

Here we report the first in-depth study of HLA presented neoantigens in esophageal adenocarcinoma, investigating both direct observation and predicted neoantigens from seven patients.

The mutational landscape of these EAC patients described by WES is consistent with previous characterizations of high mutational burden [13], mutational signatures with high proportions of C>T substitutions and evidence of chromosomal instability [4,5,20]. Gene expression analysis estimating the populations of infiltrating lymphocytes indicated that mutations yielding neoantigens may be detectable. However, only one neoantigen could be identified following direct examination of neoantigens using mass spectrometry-based proteomics to identify HLA bound peptides eluted from tumor tissues. This is consistent with previous attempts at direct neoantigen identification across multiple cancer types [6,21,22].

Although we were unable to validate the functionality of this observed neoantigen due to unavailability of PBMCs for this individual, the observed neoantigen had an optimum length and binding motif for one of the patients HLA molecules. The G>R mutation changes this peptide from a wildtype peptide that would not be expected to bind to HLA and therefore not be presented, to a peptide that can bind and be presented. Hence, we believe that this is likely to be a true neoantigen, although we are unable to confirm if it is also immunogenic.

Evidence from checkpoint inhibitor therapy and T-cell responses to predicted neoantigens suggest that neoantigens are effective at eliciting immune responses [23–25]. Therefore, for another patient for which there was available PBMCs we used the mutational and gene expression information to generate ranked lists of predicted neoantigens for each HLA-A, B and HLA-DRB1 allotype [18,19]. We tested nine of the top ranked predicted neoantigens and their wildtype equivalents for their ability to stimulate T-cells in an IFN-□ release cultured ELISpot assay and found a single high responding neoantigen (>3,000 spots/million cells, the wildtype peptide did not respond.) As with our attempts at direct identification, a one in nine success is comparable with previous reported attempts at predicting functional neoantigens [7]. The responding 15-mer peptide was a HLA-DRB1 predicted neoantigen, but on examination the first nine amino acids also comprised a HLA-A neoantigen for this patient.

The identification of a neoantigen containing both a HLA-I and HLA-II motif corresponds with reports of primarily CD4 responsive neoantigens even where neoantigens have been predicted as HLA-I peptides eliciting CD8 responses [24,25]. Similar observations have been reported in studies for viral pathogen specific peptides of CD4 responses where CD8 responses would be expected to predominate [26]. A feature of many neoantigen studies is the use of long peptides containing the neoantigen sequence and the reliance on cellular machinery to process these 20-25mer peptides into appropriate HLA-I or HLA-II length neoantigens [25,27]. Without further deconvolution, such as pre-enrichment for CD8 T-cells prior to ELISpot or single cell RNAseq TCR analysis, it is unclear what peptide processing has occurred and what immune response is being observed [28]. Here we used the specific peptides, but there still remains uncertainty about whether this a combined CD4/CD8 response or only CD4 response. Likewise the reasons for reports of predominantly CD4 responses, especially to HLA-I ligands, remains unclear.

The main limitation of our study was the availability of PBMCs for validation of putative neoantigens. Our identification of a functional neoantigen in one patient suggests that we would be able to identify others across the cohort if we were able to test them.

Overall, this study confirms that either direct observation or prediction of functional neoantigens is rare with existing methodologies, and thus further work is required to increase the frequency of successful identification [28]. However, our study also demonstrates that identified neoantigens can yield strong immune responses in functional assays, highlighting the potential for the development of neoantigen based T-cells vaccines and expanding the treatment options for a cancer with low survivability.

## Materials and Methods

### Ethics statement

Informed written consent was provided for participation by all individuals. Ethical approval for this study was granted by the Proportionate Review Sub-Committee of the North East - Newcastle & North Tyneside 1 Research Ethics Committee (Reference 18/NE/0234). This study was approved by the University of Southampton Research Ethics Committee.

### Tissue preparation

Seven subjects diagnosed with esophageal adenocarcinoma were recruited to the study (see Table 1 for clinical characteristics). tumors were excised from resected esophageal tissue post-operatively by pathologists and processed either for histological evaluation of tumor type and stage, or snap frozen at −80°C. Whole blood samples were obtained, and PBMCs were isolated by density gradient centrifugation over Lymphoprep prior to storage at −80°C.

### HLA typing

HLA typing was performed by Next Generation Sequencing by the NHS Blood and Transplant Histocompatibility and Immunogenetics Laboratory, Colindale, UK.

### DNA and RNA extraction

DNA and RNA were extracted from tumor tissue that had been obtained fresh and immediately snap frozen in liquid nitrogen. Ten to twenty 10 μm cryosections were used for nucleic acid extraction using the automated Maxwell^®^ RSC instrument (Promega) with the appropriate sample kit and according to the manufacturer’s instructions: Maxwell RSC Tissue DNA tissue kit and Maxwell RSC simplyRNA tissue kit, respectively. Similarly, DNA was extracted from snap frozen normal adjacent esophagus tissue as described above. DNA and RNA were quantified using Qubit fluorometric quantitation assay (ThermoFisher Scientific) according to the manufacturer’s instructions. RNA quality was assessed using the Agilent 2100 Bioanalyzer generating an RNA integrity number (RIN; Agilent Technologies UK Ltd.).

### Whole exome sequencing

The tumor and normal adjacent samples were prepared using SureSelect Human All Exon V7 library (Agilent, Santa Clara USA). 100 bp paired end reads sequencing was performed using the Illumina NovaSeq 6000 system by Edinburgh Genomics (Edinburgh, UK) providing ~100X depth. Reads were aligned to the 1000 genomes project version of the human genome reference sequence (GRCh38/hg38) using the Burrows-Wheeler Aligner (BWA; version 0.7.17) using the default parameters with the addition of using soft clipping for supplementary alignments. Following GATK Best Practices, aligned reads were merged [29], queryname sorted, de-duplicated and position sorted [30] prior to base quality score recalibration [31].

### Somatic variant calling

Somatic variant calling was performed using three variant callers: Mutect2 (version 4.1.2.0) [32], Varscan (version 2.4.3) [33], and Strelka (version 2.9.2) [34]. For Mutect2, a panel of normals was created using 40 samples (20 male and 20 female) from the GBR dataset. Variants were combined using gatk GenomeAnalysisTK (version 3.8-1) with a priority order of Mutect2, Varscan, Strelka. Variants were then left aligned and trimmed, and multi-allelic variants split [35]. Hard filtering of variants was performed such that only variants that had a variant allele fraction > 5%, a total coverage > 20 and variant allele coverage > 5 were kept. Filtered variants were annotated using VEP (version 97) [36] and with their read counts (https://github.com/genome/bam-readcount) to generate the final filtered and annotated variant call files (VCF).

### RNA sequencing

Samples were prepared TruSeq unstranded mRNA library (Illumina, San Diego, USA) and paired sequencing was performed using the Illumina NovaSeq 6000 system by Edinburgh Genomics (Edinburgh, UK). Raw reads were pre-processed to using fastp (version 0.20.0) [37]. Filtered reads were aligned to the 1000 genomes project version of the human genome reference sequence (GRCh38/hg38 using hisat2 (version 2.1.0) [38], merged and then transcripts assembled and gene expression estimated with stringtie2 (version 1.3.5) [39] using reference guided assembly.

### Mutanome generation

The annotated and filtered VCFs were processed using Variant Effect Predictor (version 97) [36] plugin ProteinSeqs to derive the amino acid sequences arising from missense mutations for each sample for use in immunopeptide analyses.

### Neoantigen prediction

Variant call files were prepared for the pvacseq neoantigen prediction pipeline (version 1.5.1) [18,19] by adding tumor and normal DNA coverage, and tumor transcript and gene expression estimates using vatools (version 4.1.0) (http://www.vatools.org/). Variant call files of phased proximal variants were also created for use with the pipeline [40]. Prediction of neoantigens arising from somatic variants was then performed using pvacseq with the patient HLA allotypes to predict 8-11mer peptides for class I HLA and 15-mer peptides for class II HLA-DRB allotypes. Eight binding algorithms were used for class I predictions (MHCflurry, MHCnuggetsI, NNalign, NetMHC, PickPocket, SMM, SMMPMBEC, SMMalign) and four for class II predictions (MHCnuggetsII, NetMHCIIpan, NNalign, SMMalign). Unfiltered outputs were post-processed in R [41] and split into individual tables for each peptide length and HLA allotype for each patient, and each table was then ranked according to the pvacseq score, where:

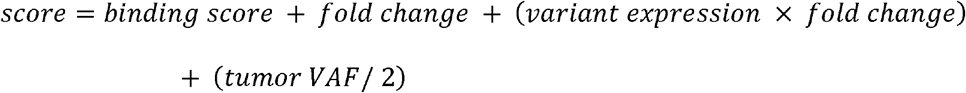

Here *binding score* is 1/median neoantigen binding affinity, *fold change* is the difference in median binding affinity between neoantigen and wildtype peptide (agretopicity). The ranked tables of predicted neoantigens were then used as described in the results.

### Immunopeptidomics

Snap frozen tissue samples were briefly thawed and weighed prior to 30s of mechanical homogenization (Fisher, using disposable probes) in 4 mL lysis buffer (0.02M Tris, 0.5% (w/v) IGEPAL, 0.25% (w/v) sodium deoxycholate, 0.15mM NaCl, 1mM EDTA, 0.2mM iodoacetamide supplemented with EDTA-free protease inhibitor mix). Homogenates were clarified for 10 min at 2,000g, 4°C and then for a further 60 min at 13,500g, 4°C. 2 mg of anti-MHC-I mouse monoclonal antibodies (W6/32) covalently conjugated to Protein A sepharose (Repligen) using DMP as previously described [42,43] were added to the clarified supernatants and incubated with constant agitation for 2 h at 4°C. The captured MHC-I/□~2~m/immunopeptide complex on the beads was washed sequentially with 10 column volumes of low (isotonic, 0.15M NaCl) and high (hypertonic, 0.4M NaCl) TBS washes prior to elution in 10% acetic acid and dried under vacuum. The MHC-I-depleted lysate was then incubated with anti-MHC-II mouse monoclonal antibodies (IVA12) and MHC-II bound peptides were captured and eluted in the same conditions.

Immunopeptides were separated from MHC-I/□~2~m or MHC-II heavy chain using offline HPLC on a C18 reverse phase column. Briefly, dried immunoprecipitates were reconstituted in buffer (1% acetonitrile,0.1% TFA) and applied to a 10cm RP-18e chromolith column using an Ultimate 3000 HPLC equipped with UV monitor. Immunopeptides were then eluted using a 15 min 0-40% linear acetonitrile gradient at a flow rate of 1 mL/min.

HLA peptides were separated by an Ultimate 3000 RSLC nano system (Thermo Scientific) using a PepMap C18 EASY-Spray LC column, 2 μm particle size, 75 μm x 75 cm column (Thermo Scientific) in buffer A (0.1% Formic acid) and coupled on-line to an Orbitrap Fusion Tribrid Mass Spectrometer (Thermo Fisher Scientific,UK) with a nano-electrospray ion source. Peptides were eluted with a linear gradient of 3%-30% buffer B (Acetonitrile and 0.1% Formic acid) at a flow rate of 300 nL/min over 110 minutes. Full scans were acquired in the Orbitrap analyser using the Top Speed data dependent mode, preforming a MS scan every 3 second cycle, followed by higher energy collision-induced dissociation (HCD) MS/MS scans. MS spectra were acquired at resolution of 120,000 at 300 m/z, RF lens 60% and an automatic gain control (AGC) ion target value of 4.0e5 for a maximum of 100 ms. MS/MS resolution was 30,000 at 100 m/z. Higher□energy collisional dissociation (HCD) fragmentation was induced at an energy setting of 28 for peptides with a charge state of 2–4, while singly charged peptides were fragmented at an energy setting of 32 at lower priority. Fragments were analysed in the Orbitrap at 30,000 resolution. Fragmented m/z values were dynamically excluded for 30 seconds.

### Proteomic data analysis

Raw spectrum files were analyzed using Peaks Studio 10.0 build 20190129 [15,44] and the data processed to generate reduced charge state and deisotoped precursor and associated product ion peak lists which were searched against the UniProt database (20,350 entries, 2020-04-07) plus the corresponding mutanome for each sample (~1,000-5,000 sequences) and contaminants list in unspecific digest mode. Parent mass error tolerance was set a 5ppm and fragment mass error tolerance at 0.03 Da. Variable modifications were set for N-term acetylation (42.01 Da), methionine oxidation (15.99 Da), carboxyamidomethylation (57.02 Da) of cysteine.

A maximum of three variable modifications per peptide was set. The false discovery rate (FDR) was estimated with decoy-fusion database searches [15] and were filtered to 1% FDR. Downstream analysis and data visualizations of the Peaks Studio identifications was performed in R using associated packages [41,45].

### Immunopeptide HLA assignment

Identified immunopeptides were assigned to their HLA allotype for each patient using motif deconvolution tools and manual inspection. For class I HLA peptides initial assignment used MixMHCp (version 2.1) [6,17] and for class II HLA peptides initial assignment used MoDec (version 1.1) [46]. Downstream analysis and data visualizations was performed in R using associated packages [41,45,47].

### Synthetic peptides

Peptides for functional T-cell assays and spectra validation were synthesised using standard solid phase Fmoc chemistry (Peptide Protein Research Ltd, Fareham, UK).

### Functional T-cell assay

PBMC (2×10^6^ per well) were stimulated in 24-well plates with peptide (individual/pool) plus recombinant IL-2 (R&D Systems Europe Ltd.) at a final concentration of 5μg/mL and 20IU/mL, respectively, and incubated at 37°C with 5% CO2; final volume was 2mL. Media containing additional IL-2 (20IU/mL) was refreshed on days 4, 6, 8 and 11 and on day 13 cells were harvested. Expanded cells (1×10^5^ cell/well) were incubated in triplicate with peptide (individual) at 5μg/mL final concentration for 22 hours at 37°C in 5% CO2; phytohemagglutinin (PHA; Sigma-Aldrich Company Ltd.) and CEFT peptide mix (JPT Peptide Technologies GmbH, Berlin, Germany), a pool of 27 peptides selected from defined HLA Class I- and II-restricted T-cell epitopes, were used as positive controls. Spot forming cells (SFC) were counted using the AID ELISpot plate reader system ELR04 and software (AID Autoimmun Diagnostika GmbH) and positivity calling for ELISpot data used the runDFR(x2) online tool (http://www.scharp.org/zoe/runDFR/). Downstream analysis and data visualizations was performed in R using associated packages [41,45].

## Data availability

The mass spectrometry proteomics data have been deposited to the ProteomeXchange Consortium via the PRIDE [48] partner repository with the dataset identifier PXD031108 and 10.6019/PXD031108.

Reviewer account details: Username: Password: g598EYGq

Whole exome and RNA sequencing data has been deposited at the European Genome-phenome Archive (EGA), which is hosted by the EBI and the CRG, under accession number EGAS00001005957.

The authors declare that all the other data supporting the finding of this study are available within the article and its supplementary information files and from the corresponding author on reasonable request.

## Acknowledgements

This study was supported by a Cancer Research UK Centres Network Accelerator Award Grant (C328/A21998). Instrumentation in the Centre for Proteomic Research is supported by the BBSRC [BM/M012387/1]

