## supporting information for "Identification of neoantigens in esophageal adenocarcinoma"

### Extended patient information

**S1 Table: Extended summary of individuals with oesophageal adenocarcinoma**

| Donor | ICGC Donor | Age | Sex | Tumour location | Stage | Treatment | Tumour Mut/Mb | Healthy Mut/Mb | Tumour somatic variants | Healthy somatic varaints | Obs.HLA-I peps | Obs.HLA-II peps | HLA-I | HLA-II |
|---|---|---|---|---|---|---|---|---|---|---|---|---|---|---|
| EN-181-11 | DO234382 | 64 | Male | Gastro-Oesophageal junction (Siewert II) | 2 | Surgery | 142 | 40 | 5,049 | 1,430 | 5,332 | 2,420 | A*03:01,B*07:02,B*08:01,C*07:01,C*07:02 | DRB1*03:01,DRB1*15:01,DQB1*02:01, DQB1*06:02,DPB1*01:01,DPB1*04:01 |
| EN-216-11 | DO50307 | 74 | Male | Lower Oesophagus | 2 | Surgery | 101 | 41 | 3,595 | 1,468 | 7,250 | 4,531 | A*02:01,A*29:02,B*15:01,B*18:01,C*03:04, C*05:01 | DRB1*03:01,DRB1*04:01,DQB1*02:01, DQB1*03:02,DPB1*02:02,DPB1*04:01 |
| EN-430-11 | | 78 | Male | Lower Oesophagus | 3 | Chemotherapy + Surgery | 124 | 41 | 4,430 | 1,446 | 1,411 | 0 | A*01:01,A*11:01,B*08:01,B*35:01,C*04:01, C*07:01 | DRB1*01:01,DRB1*10:01,DQB1*05:01, DPB1*04:01 |
| EN-454-11 | DO50387 | 81 | Male | Gastro-Oesophageal junction (Siewert II) | 3 | Surgery | 438 | 40 | 15,651 | 1,416 | 7,389 | 1,553 | A*01:01,A*24:01,B*08:01,B*15:01,C*03:03, C*07:01 | DRB1*03:01,DRB1*13:02,DQB1*02:01, DQB1*06:04,DPB1*02:01,DPB1*04:01 |
| EN-489-12 | | 62 | Male | Lower Oesophagus | 2 | Chemotherapy + Surgery | 120 | 42 | 4,271 | 1,495 | 5,430 | 1,560 | A*23:01,A*33:01,B*14:02,B*49:01,C*07:01, C*0:02 | DRB1*08:03,DRB1*15:01,DQB1*03:01, DQB1*06:02,DPB1*03:01,DPB1*04:01 |
| EN-711-16 | | 68 | Male | Lower Oesophagus | 3 | Chemotherapy + Surgery | 115 | 40 | 4,097 | 1,413 | 2,819 | 71 | A*01:01,A*23:01,B*08:01,B*44:03,C*04:01, C*07:01 | DRB1*03:01,DRB1*07:01,DQB1*02:01, DQB1*02:02,DPB1*01:01,DPB1*02:01 |
| EN-716-11 | | 66 | Male | Lower Oesophagus | 3 | Chemotherapy + Surgery | 136 | 39 | 4,865 | 1,397 | 1,521 | 248 | A*02:01,B*44:02,B*58:01,C*05:01,C*07:18 | DRB1*04:01,DRB1*13:02,DQB1*03:01, DQB1*06:09,DPB1*01:01,DPB1*04:01 |

Age: Age at diagnoses. Stage: Tumour stage. Treatment: Treatment modality. Obs.HLA-I peps: Observed HLA-I peptides. Obs.HLA-II peps: Observed HLA-II peptides. HLA-I: HLA-I allotypes. HLA-II: HLA-II allotypes.

**HLA motifs**

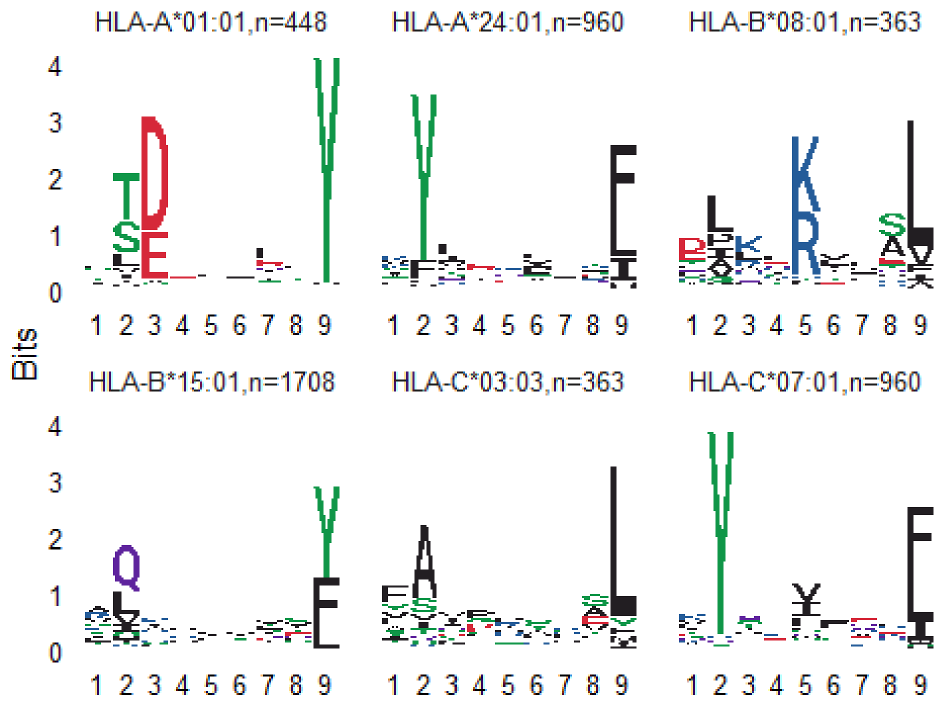


***S1 Figure: EN-454-11 HLA-I 9mer motifs Eluted peptides clustered using MixMHC and then HLA manually assigned.***

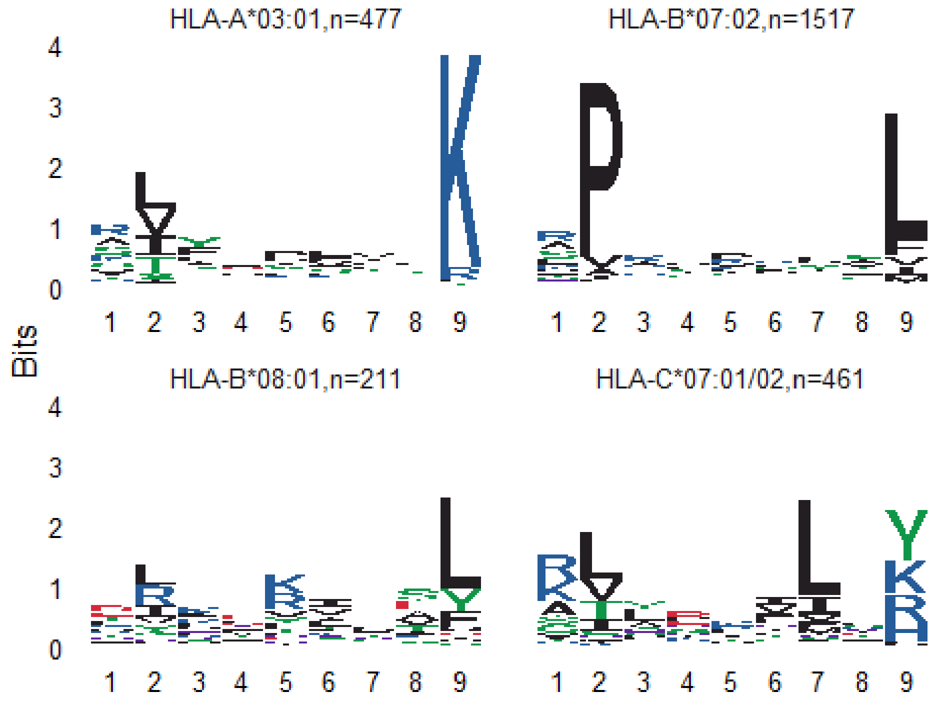


***S2 Figure: EN-181-11 HLA-I 9mer motifs Eluted peptides clustered using MixMHC and then HLA manually assigned.***

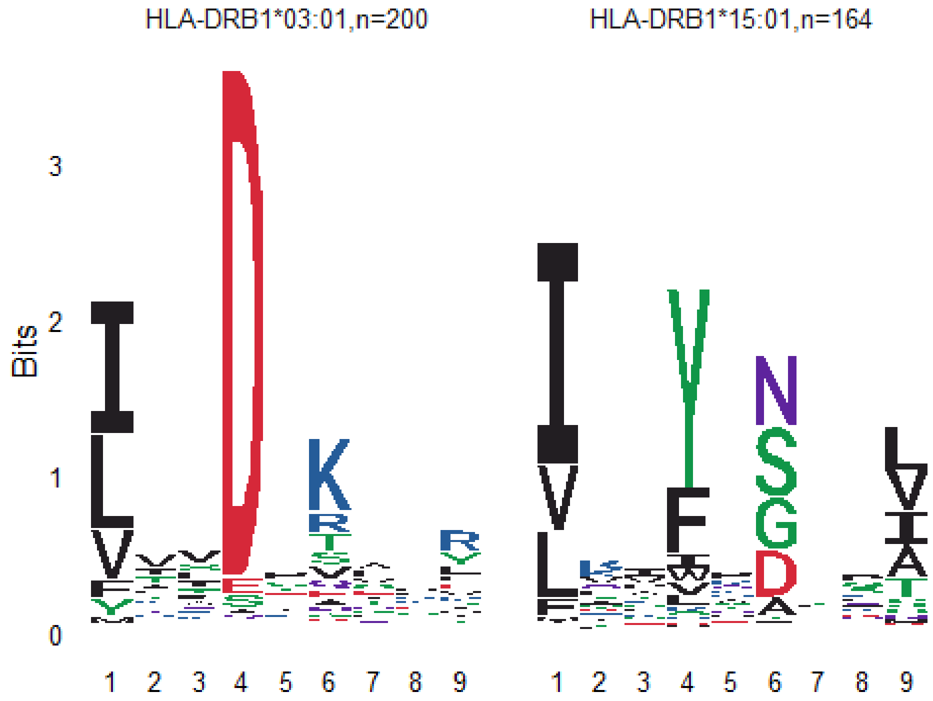


***S3 Figure: EN-181-11 HLA-II DRB1 core motifs Eluted peptides clustered using MixMHC and then HLA manually assigned.***
